# The Bayesian Phylogenetic Bootstrap, Application to Short Trees and Branches

**DOI:** 10.1101/2024.06.22.600199

**Authors:** Frédéric Lemoine, Olivier Gascuel

## Abstract

Felsenstein’s bootstrap is the most commonly used method to measure branch support in phylogenetics. Current sequencing technologies can result in massive sampling of taxa (e.g. SARS-CoV-2). In this case, the sequences are very similar, the trees are short, and the branches correspond to a small number of mutations (possibly 0). Nevertheless, these trees contain a strong signal, with unresolved parts but a low rate of false branches. With such data, Felsenstein’s bootstrap is not satisfactory. Due to the frequentist nature of bootstrap sampling, the expected support of a branch corresponding to a single mutation is ∼63%, even though it is highly likely to be correct. Here we propose a Bayesian version of the phylogenetic bootstrap in which sites are assigned uninformative prior probabilities. The branch support can then be interpreted as a posterior probability. We do not view the alignment as a small subsample of a large sample of sites, but rather as containing all available information (e.g., as with complete viral genomes, which are becoming routine). We give formulas for expected supports under the assumption of perfect phylogeny, in both the frequentist and Bayesian frameworks, where a branch corresponding to a single mutation now has an expected support of ∼90%. Simulations show that these theoretical results are robust to realistic data. Analyses on low-homoplasy viral and non-viral datasets show that Bayesian bootstrap support is easier to interpret, with high supports for branches very likely to be correct. As homoplasy increases, the two supports become closer and strongly correlated.

## Introduction

The phylogenetic bootstrap was proposed by Joseph Felsenstein to estimate the branch supports of trees inferred from multiple sequence alignments (Felsenstein 1985). Given a multiple sequence alignment (MSA) and a reference tree inferred from it, the principle is to sample with replacement the sites of the original alignment to obtain pseudo-alignments of the same length, infer pseudo-trees from the pseudo-alignments, and measure the support of each branch in the reference tree as the proportion of pseudo-trees containing that branch. The usefulness and simplicity of this method made it extremely popular. Felsenstein’s 1985 article is among the top 100 most cited scientific papers of all time (Van Noorden et al. 2014).

The phylogenetic bootstrap is an application to MSA and trees, of the statistical bootstrap, which is commonly used to study the distribution of numerical estimates (Efron 1979; Efron and Tibshirani 1993). In this context, the sampling with replacement procedure has a frequentist basis and mimics the sampling of observations (or statistical units) in a large set of homogeneous and independent observations. The relevance and limits of these homogeneity and independence assumptions have been questioned on a biological basis (Sanderson 1995) when applied to the phylogenetic bootstrap, where MSA sites are considered as statistical units. The complexity of phylogenetic tree spaces and sequence evolution models (compared to simpler numerical spaces and estimates) makes statistical interpretation of bootstrap supports difficult (Holmes 2003). This issue was the subject of much debate in the 1990s (Hillis and Bull 1993; Felsenstein and Kishino 1993; Berry and Gascuel 1996). Several methods (Zharkikh and Li 1995; Efron et al. 1996; Susko 2009, 2010) have been proposed to correct bootstrap supports to better agree with standard ideas of confidence levels and hypothesis testing. However, bootstrap correction methods are limited to relatively small data sets, and the original method is still highly used (∼2,000 citations in 2023). As stated by Soltis and Soltis (2003): “consensus has been reached among practitioners, if not among statisticians and theoreticians”, and “many systematists have adopted Hillis and Bull’s “70%” value as an indication of support”. This 70% threshold indicates the very conservative nature of the standard bootstrap. The bootstrap is also computationally heavy, but it is easily parallelized and fast algorithms have been designed (Stamatakis et al. 2008; Minh et al. 2013).

Alternatives to the Felsenstein bootstrap are local supports (aLRT; Anisimova et al. 2006, Guindon et al. 2010), which are very fast to compute, but can be too local and support branches that should not be supported from a global perspective (Gascuel and Lemoine 2024). Bayesian posterior probabilities are also widely used, but they tend to be liberal (Suzuki et al. 2003), are difficult to apply to large datasets, and are sensitive to model misspecification (Huelsenbeck and Rannala 2004). The aBayes support (Anisimova et al. 2011) is a local approximation of the posterior branch probability; it is fast to compute, and has recently been reported for its accuracy on virus datasets (Wertheim et al. 2022). A common practice among phylogeneticists is to compute both the bootstrap support (conservative) and the Bayesian posterior (liberal) to obtain a fair view of the evidence supporting the branches, given the data and priors (Douady et al. 2003). Recently, several machine-learning approaches have been proposed (Wiegert et al. 2024; Ecker et al. 2024) to combine various indicators and fast supports (e.g., aLRT statistic, parsimony-based bootstrap).

The Felsenstein bootstrap and most of the approaches mentioned above face the problem of “rogue” taxa (Sanderson and Shaffer 2002; Soltis and Soltis 2003), which are inevitable in large datasets with many taxa. These taxa are unstable within bootstrap trees, and the presence of a single rogue taxon in a dataset makes the bootstrap support very low when a branch is found almost identically in bootstrap trees, except for one or a few misplaced rogue taxa. To address this problem, we proposed the Transfer Bootstrap Expectation (TBE, Lemoine et al. 2018; Zaharias et al. 2023), in which we measure the degree of presence of reference branches in bootstrap trees using a quasi-continuous index (as opposed to the binary index used in the standard bootstrap). The support of a reference branch corresponds to the average number of taxa that must be transferred (or deleted) to find that branch in the bootstrap trees. TBE has been shown to yield higher and more informative supports than the standard approach, while inducing a very low number of falsely supported branches.

The Felsenstein bootstrap is also problematic with short trees and branches supported by only a few mutations (Wertheim et al. 2022). This type of data is becoming increasingly common with the massive sequencing of many species, especially viruses such as SARS-CoV-2, Ebola virus (EBOV), Zika virus (ZIKV), Dengue virus (DENV) and many others (Dudas and Bedford 2019). However, viruses are not the only organisms concerned. Studies of closely related species and intraspecific analyses also produce this type of data and phylogeny, as well as single-cell data, which is increasingly used in cancer and developmental biology (Jahn et al. 2016). Under these conditions, we are close to perfect (or Hennigian) phylogeny assumptions (Felsenstein 1973; Gusfield 1997; Ramazzotti et al. 2021; Wertheim et al. 2022). The level of homoplasy is low or even zero at some sites, and the sites contradict each other little or not at all. Contradictions between sites (e.g., one site supports cherry {A, B} with a mutation separating taxa A and B from all other taxa, while another contradictory site supports cherry {A, C}) can be the result of random neutral mutations or, for example, caused by recombination and horizontal transfer. Under these near-perfect conditions, Wertheim et al. (2022) show that phylogenies are poorly resolved, but that inferred branches have a high probability of being correct. However, theoretical analyses (Felsenstein 1985) show that the expected bootstrap support for a branch having undergone a single mutation is ∼0.632 (63%). This value is well known in bootstrap theory (Efron and Tibshirani 1997), and corresponds to the probability that a given site (e.g. carrying a mutation supporting a given clade) will be present at least once in a bootstrap sample randomly drawn with replacement. This is a clear shortcoming of the standard bootstrap when the data is near-perfect, and (Wertheim et al. 2022) recommends against bootstrap resampling when substitutions are sparse.

In fact, in the standard frequentist bootstrap, certain sites are removed (with probability 1 - 0.632 ≈ 37%) and part of the original information is lost, resulting in bootstrap samples that are less informative than the original sample. This becomes problematic with small samples and when the information is poor, for example with virus data, where a large number of sites are invariant. From a biological point of view, this resampling method can also be questioned, as it implicitly assumes that the original sample is randomly taken from a larger sample, typically a genome. However, in the case of viruses (e.g., SARS-CoV-2, EBOV …) the data commonly corresponds to the entire genome, and we cannot think of it as a sample from a larger dataset. Sampling with replacement creates “pseudo-genomes” that are poorer in information than the original data, with only ∼63% of the sites present in the complete genome. The situation is similar when we are studying the phylogeny of a particular gene or genome segment (e.g., in influenza, which is prone to reassortment), the history of which may be different from that of the genomes and species/strains from which it was extracted; again, the information that is available in its entirety is in the input MSA.

In this paper, we propose a Bayesian approach to phylogenetic bootstrapping in which all sites remain present in the bootstrap samples. This approach is derived from the Bayesian bootstrap proposed by Rubin (1981) and others (Shao and Tu 1995) in the context of numerical data and estimations. The principle is to weight the *n* observations of the original sample, rather than drawing them with replacement. Each observation *i* thus has a weight *w*_*i*_ drawn from a continuous non-informative prior distribution that is identical and independent for all observations (or statistical units). Rubin suggests using a Dirichlet distribution, but other distributions have been studied (Lo 1991). Theoretical results show that, under certain conditions, the distribution obtained for the estimated parameters (e.g., the mean or variance of a normal distribution) converges to the posterior of those parameters when their priors are non-informative (Lo 1991). The Bayesian bootstrap is a relevant alternative to the frequentist bootstrap, which is smoother and generally more suitable for small samples (Shao and Tu 1995). In the following, we introduce and study the application of this approach to phylogenetics. We provide the expected values for standard frequentist and Bayesian bootstrap supports under the assumption of perfect phylogeny. We compare the results of the two approaches on simulated data that gradually deviate from perfect phylogeny. We analyze real datasets of viruses (EBOV, SARS-CoV-2, RVFV) as well as a plant dataset (*Psiguria* genus, *Cucubirtacea*) with low homoplasy. The results show that for this type of data with low homoplasy, Bayesian bootstrap supports bring some gain in accuracy, are easier to interpret, with high supports for branches that are essentially correct. As homoplasy increases, the two supports become closer and strongly correlated.

## Methods and theoretical analyses

The data is a multiple sequence alignment (MSA; DNA or protein) containing *n* sites and *s* sequences. The goal is to construct a phylogenetic tree from this MSA and measure the support of its branches. To simplify the presentation, we will mainly use the maximum likelihood ML framework, but the methods presented can easily be adapted to other approaches.

The length of branches in a phylogeny is expressed as the expected number of mutations per site. If an internal branch has a length of zero, this essentially means that it is not supported by any of the mutations present in the input MSA and that we are in the presence of a polytomy. Such a case is common in short trees, especially in viral phylogenies, and requires special attention in the bootstrap framework. In the following, we discuss the treatment of (near) zero branches, and then examine the support of branches carrying one or more mutations (i.e., of non-zero length), first in the framework of the standard frequentist phylogenetic bootstrap, and then in the framework of the Bayesian bootstrap. Finally, we present several criteria for evaluating and comparing different types of branch support.

### Polytomies, (near) zero branches, and bootstrap support

If an inferred internal branch has length (near) zero, this means that the inference program did not find any direct or multiple mutations along that branch in the MSA, which should be collapsed to create a multifurcation (or polytomy). Parsimony programs can produce polytomies and branches of strictly zero length because they count mutations that occur along tree branches, but ML programs do not, due to numerical optimization constraints. ML programs define a minimum branch length, and if a length estimated by ML reaches the minimum, the corresponding branch should be considered null. For example, the default minimum branch length in IQTREE (Minh et al. 2020) is “the smaller of 10^−6^ and 0.1/alignment_length”, and it is recommended to use the ‘czb’ option, which “Collapses near zero branches, so that the final tree may be multifurcating. This is useful for bootstrapping in the presence of polytomy to reduce bootstrap supports of short branches”. Similarly, RAxML-NG (Kozlov et al. 2019) output contains the “bestTreeCollapsed”, where near-zero branches are collapsed (default minimum length is of 10^−6^).

With virus and low homoplasy datasets, near-zero branches are common, in particular due to the presence of duplicated sequences, but not only (e.g., Fig. 1). In bootstrap trees, this phenomenon is further amplified, since a branch having undergone a unique mutation in the original MSA may be not inferred anymore, if the corresponding site is not drawn in the bootstrap MSA. Moreover, the primary output of ML inference programs is a binary tree. If a branch has near-zero length, this actually corresponds to a polytomy whose resolution through the given branch does not reflect any signal in the input data. However, this resolution is generally non-random. For example, RaxML-NG and PhyML (Guindon et al. 2010) tend to produce caterpillars in unresolved parts of the tree, and the order of the sequences in the MSA play a part in the resolution of polytomies. The effect of these hidden determinisms, which vary from program to program, can cause some near-zero branches to have high bootstrap support, a problem that is remedied by the collapse of near-zero branches and the use of multifurcating trees.

**Figure 1:**
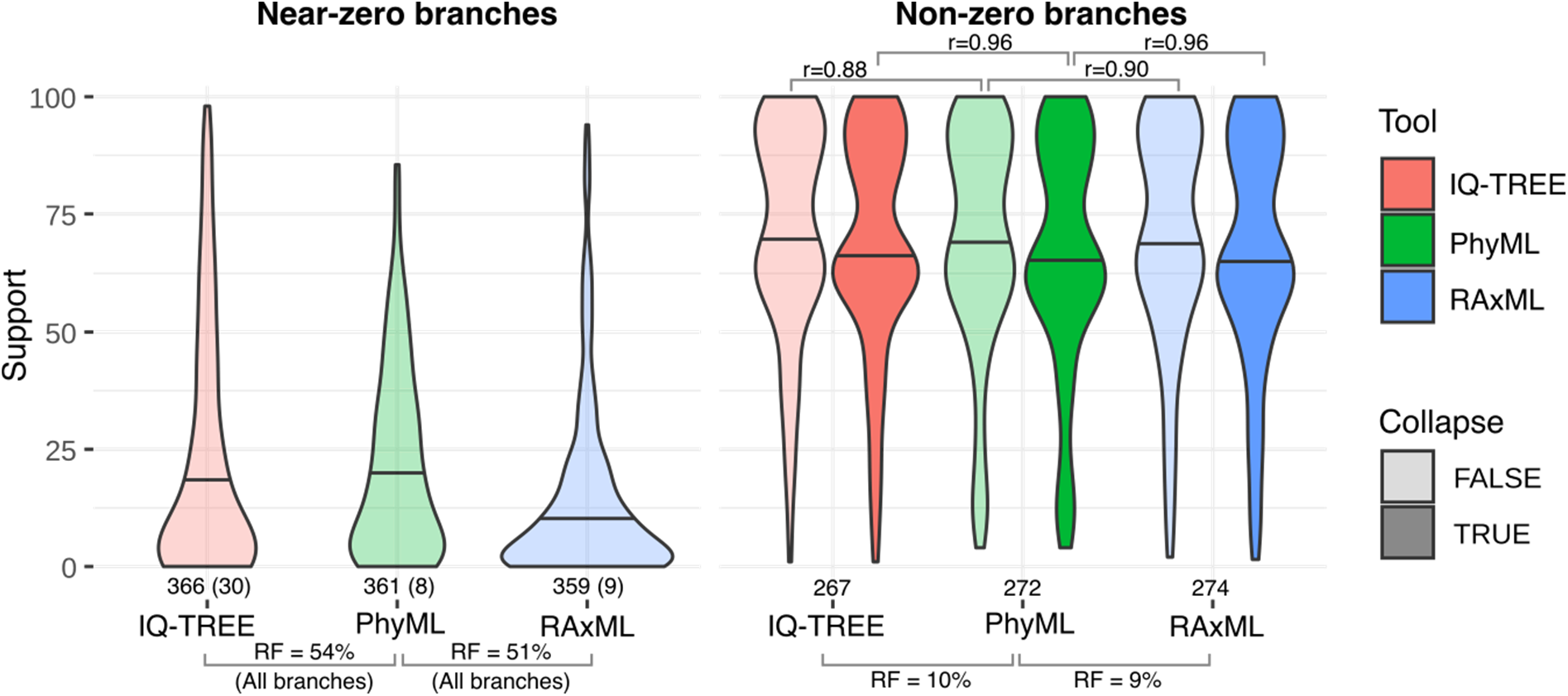
Distribution of frequentist bootstrap supports with/without collapse of near-zero branches, using Ebola virus full-genome data. The MSA (extracted from Dudas et al. 2017) contains 636 unique sequences. **(a)** Distribution of supports of near-zero branches without collapse (neither the reference tree nor the bootstrap trees). We provide: (i) the number of such branches, e.g. 366 for IQTREE, out of a total of 633 internal branches; (ii) the number (in parentheses) of such branches with support ≥70%, e.g. 30 for IQTREE; the Robinson and Foulds (RF) topological distance between non-collapsed trees, e.g. 54% between the reference trees inferred by IQTREE and PHYML. **(b)** Distribution of branch supports of non-zero branches. Light coloring: near-zero branches in bootstrap trees are not collapsed. Dark coloring: near-zero branches in bootstrap trees are collapsed. We provide: (i) the number of non-zero internal branches in the reference tree, e.g. 267 for IQTREE; (ii) the RF distance between the collapsed reference trees, e.g. 10% for IQTREE and PhyML; (iii) the correlation of the supports for the inferred branches shared by the two programs with the two collapse options, e.g. 0.96 between IQTREE and PhyML when both reference and bootstrap trees are collapsed, and 0.88 without collapse when all shared (near-zero and non-zero) branches are considered.

We illustrate all this using an Ebola virus (EBOV) full-genome dataset extracted from (Dudas et al. 2017; see below and Sup. Mat. for details). The original MSA contains 1,610 sequences and 18,996 sites, from which we randomly subsampled 800 sequences for computational reasons. We then removed duplicate sequences and obtained an MSA with 636 unique sequences. We ran IQTREE, RaxML-NG, and PhyML on this MSA using GTR+G4 and 200 bootstrap replicates with two options: (i) no collapse of any branches, and (ii) collapse of near-zero internal branches in both the reference and bootstrap trees (e.g., using the ‘czb’ option in IQTREE with default minimum length; see Sup. Mat. for details). We compared the results of the different programs and options. The results are shown in Figure 1.

The number of near-zero internal branches is high for the three programs (>55%), even though all duplicates have been removed. Non-collapsed reference trees inferred by two different programs have high topological distances (RF >50%), as expected since near-zero branches are inferred “by chance”, and some near-zero branches have high bootstrap support (e.g., for IQTREE, 30 out of 356 near-zero branches have support ≥70%). The fraction of common branches shared by two programs is small (<50%), and on these branches the correlation between supports is not so strong (e.g., r = 0.88 for IQTREE and PhyML).

Collapsing the near-zero branches and using multifurcating trees reveals the signal in the data, and then the results of the three programs become very close. Topological distances are much smaller (∼10%), most branches are shared (∼90%), and the correlation of supports between shared branches is very high (r = 0.96, although bootstrap MSAs differ). Another notable effect is that the support of (non-zero) branches is slightly lower; this is explained by the collapse of bootstrap trees that no longer contain certain branches of the reference tree inferred by chance using bootstrap MSAs.

The collapse of near-zero branches and the use of multifurcating trees is therefore an important step in applying the standard frequentist bootstrap to virus and low-homoplasy sequences in order to extract the phylogenetic signal that is actually present in the data and to obtain reproducible results. We will see that it is also important for the Bayesian bootstrap on the same type of data, and our simulation results (see below) confirm this observation. For more divergent sequences with a high number of mutations, the importance of this trimming decreases, but it should still be considered if the reference tree has near-zero branches.

### Expected supports with the standard frequentist bootstrap

Here we analyze the expected supports with the standard frequentist bootstrap, for branches of the reference tree whose length corresponds to at least 1 mutation. In the theoretical analysis that follows, we assume that the sequence data are close to the perfect phylogeny hypothesis (Felsenstein 1973; Gusfield 1997). This implies that no or few convergent, parallel, and revertant mutations are observed at a single site. It also implies that the different sites in the alignment contain no or few conflicting signals. The hypothesis of a strictly perfect phylogeny is rarely verified, but many data sets come close, especially viral data and those with a low number of mutations and closely related sequences (Ramazzotti et al. 2021; Wertheim et al. 2022). This is the case, for example, with the EBOV and SARS-Cov-2 datasets, which we shall study in the next section. With such data, the estimated branch lengths clearly represent mutation numbers, which are generally equal to or close to 0 (Fig. 2).

**Figure 2:**
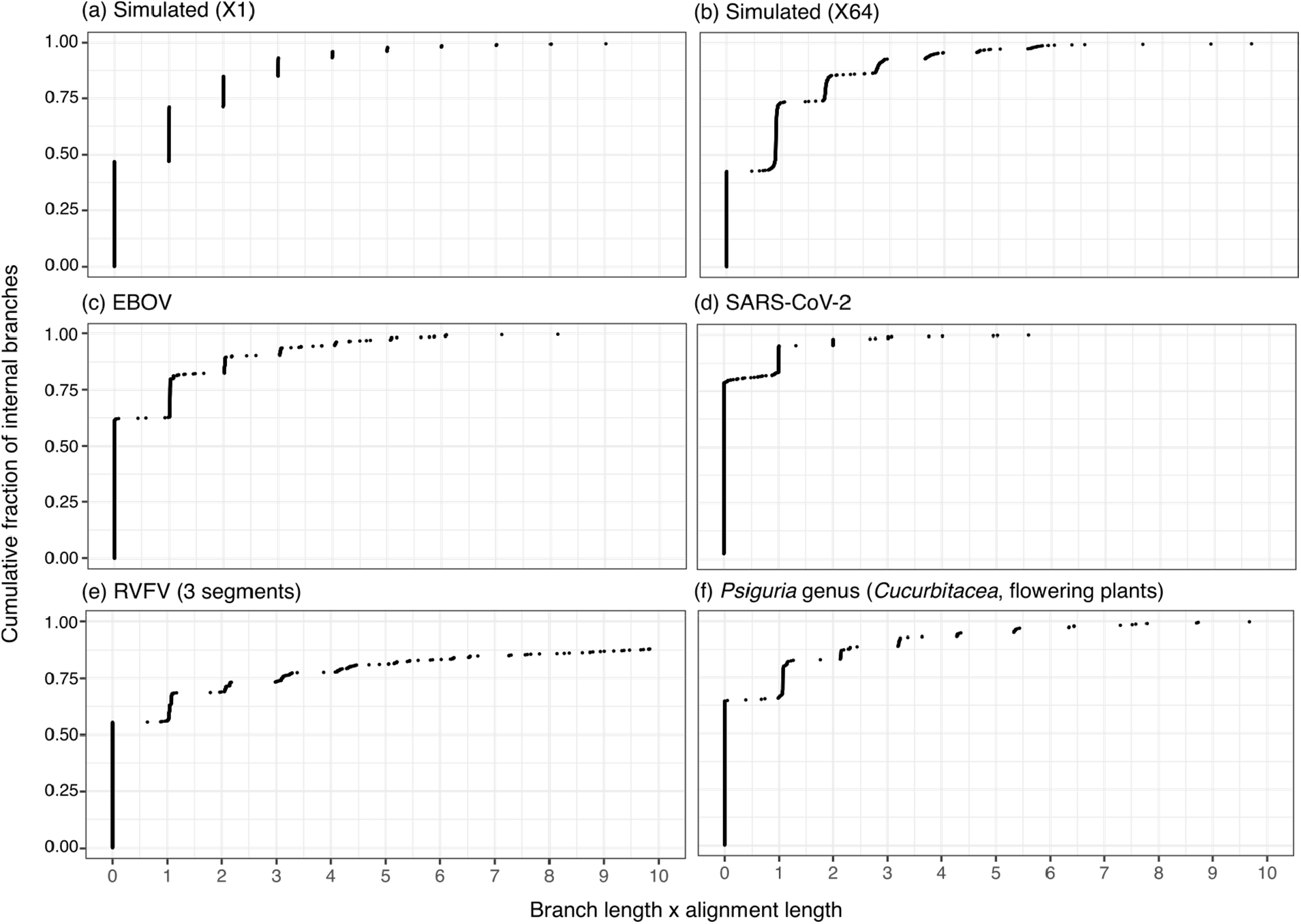
Cumulative distribution of internal branch length estimates. Branch lengths (X-axis) are expressed in number of substitutions (i.e., standard branch length multiplied by MSA length). Y-axis: cumulative fraction of internal branches. Panel **(a)**: Simulated data in X1 (EBOV-like, low homoplasy) condition. Panel **(b)**: Simulated data in X64 (high-homoplasy) condition. Panel **(c)**: EBOV data (Dudas et al. 2017). Panel **(d)**: SARS-CoV-2 data (Zhukova et al. 2021). Panel **(e)**: Three datasets of RVFV virus, segments M, L, V (combined distribution of the 3). Panel **(f)**: *Psiguria* genus (*Cucubirtacea*, flowering plants) low-homoplasy dataset. See text and Supp. Table 1 for details.

The perfect phylogeny framework was used by Felsenstein (1985) to study the relationship between the length (in number of mutations) and the bootstrap support of tree branches. We follow the same path here, starting by analyzing the case where a branch of the reference tree is supported by 1 mutation, and thus by a single site within the MSA given as input. In a bootstrap sample, the probability of this site being absent is equal to (1 −1/ *n*)^*n*^ ; in this case, the corresponding branch in the bootstrap tree will not be inferred or will have an estimated length very close to zero. Switching to the logarithm and assuming *n* is large, we can easily show that the limiting value of (1 − 1 /*n*)^*n*^ is *e*^−1^. The probability that the site is present at least 1 time and that a bootstrap tree contains and supports this branch is therefore equal to 1− *e*^−1^ ≈ 0.632. This value is well known in bootstrap theory (Efron and Tibshirani 1997). In phylogenetics, assuming (nearly) perfect data, 0.632 represents the expected bootstrap support for a branch whose length corresponds to (approximately) 1 mutation. In the case where a branch of the original tree has a length corresponding to *k* (≪ *n*) mutations, the above values become (1 − *k/ n*)^*n*^ ≈ *e*^−*k*^ and the expected bootstrap support becomes 1− *e*^−*k*^, which is approximately 0.865, 0.950, and 0.982 for *k* = 2, 3, and 4, respectively. In all of this, however, we assume that randomly inferred near-zero branches are collapsed (not accounted for) and that bootstrap trees may be multifurcating. When using the standard version of the frequentist bootstrap with full binary trees that may contain near-zero branches, the expectation of supports is slightly higher than these theoretical values (see the results that follow).

In the following, we analyze simulated and real data with a relatively low number of mutations, but deviating from the assumption of a perfect phylogeny. On these realistic/real data, the observed supports are generally very close to the theoretical ones. The simulation results also show that branches with an estimated length of ∼1 mutation (and thus a bootstrap support of ∼0.632) or more are generally correct branches (∼90% of the cases; see also Wertheim et al. 2022). This again illustrates the conservatism of the frequentist bootstrap and the difficulty of interpreting its supports, which has been discussed many times (see the Introduction).

### The Bayesian phylogenetic bootstrap, expected supports

In the Bayesian bootstrap, the observations in the bootstrap samples are drawn with weights from an a priori continuous non-informative distribution, and the resulting estimates represent (under certain assumptions) the posterior distribution of the parameter of interest (Rubin 1981; Lo 1991). The context, however, is very different from that of Bayesian inference of trees, as implemented in MrBayes for example (Huelsenbeck and Ronquist 2001). There is no prior on the parameter value, and this is purely numerical (e.g., estimating a probability or a correlation and their variability, in Rubin 1981). As Rubin points out, “the Bayesian bootstrap is a natural Bayesian analogue of the bootstrap” and the two “are quite similar inferentially”, but “the interpretations of the resulting distributions will be different” (sampling distribution for the standard bootstrap, versus posterior distribution for the Bayesian version).

Several non-informative distributions have been proposed. The most widely used is the Dirichlet distribution of parameters (*n*; 1, 1, …1). Rubin (1981) proposes a definition of this distribution and a computation based on a set of intervals in [0, 1]. An equivalent definition is to independently draw *n* weights *w*_*i*_ (attached to each of the MSA sites *i* in a phylogenetic context) in an exponential distribution with expectation 1.0 and normalize these weights by their sum. To make it closer to the usual phylogenetic framework, we multiply these normalized weights by the number of sites, i.e., *w*_*i*_ ⟵ *w*_*i*_ × *n /*(*w*_1_ + *w*_2_ +…*w*_*n*_). The sum of the normalized weights is *n*, and the expectation of each weight is 1.0, just as in the frequentist framework, where each site has a weight drawn from a multinomial distribution with parameters (*n*; 1 /*n*, 1/ *n* … 1/ *n*). The variance and the correlation of the weights over replications are also similar in both framework (Rubin 1981). The Dirichlet distribution of the weights thus represents a continuous version of the sampling with replacement procedure, and the resulting posterior parameter distribution is smoother than its frequentist analogue (Shao and Tu 1995). In phylogenetics, a major difference is that each site is present with a non-zero weight in each of the Bayesian bootstrap samples. Thus, a signal carried by a single site remains present in all samples, but with a different weight from one sample to another. Within the maximum likelihood framework, the likelihood of a weighted MSA is calculated simply by multiplying the log-likelihood of each site by its weight. Standard software programs (e.g. RaxML-NG, IQTREE) already use this weighting system with integer weights to account for identical sites. In PhyML, which we use in our experiments, it is possible to have continuous weights, as required here.

To derive the theoretical support as a function of the branch length of the original tree, we again place ourselves in the perfect phylogeny framework. In principle, in this framework and with a perfect inference method, we should find the original phylogeny regardless of the bootstrap sample and Bayesian weights, since each site is present with a positive weight and there is no conflicting signal. If a branch *b* in the reference tree is supported by 1 mutation (i.e., has length 1 /*n*), and if the weight of the corresponding site in a bootstrap sample is *w*, then the length of *b* in the corresponding bootstrap tree will be *w/ n*. If *w* is small, the length of the bootstrap branch will be very small.

In practice, short branches must be collapsed to avoid the effects of the hidden determinisms described above (Fig. 1), numerical approximations, and also because real data are never perfect, with a certain amount of homoplasy and contradiction, which can induce branches of non-zero (but almost) length that are not supported by any mutation. In our experiments, we collapsed branches of length less than 0.1 /*n* (i.e., with an estimated number of mutations less than χ =1/10 ; this value is also used in IQTREE to define the minimum branch length). Let us place ourselves in the perfect phylogeny framework and consider a reference branch *b* of length 1 /*n* ; if the weight *w* of the single site supporting this branch is less than χ =1 /10, this branch will be inferred but collapsed and therefore not counted as recovered in the corresponding bootstrap tree. The theoretical support of the branch will therefore be equal to (1 − *p*), where *p* is the probability of having a weight *w* less than χ. For relatively large *n* (say >100), the distribution of the normalized weights (see formula above) is practically equal to an exponential law with parameter 1.0, and the distribution in the neighborhood of 0 is asymptotically close to the uniform distribution in [0, 1]; in other words, *p* ≈ χ. For example, *p* is equal to ∼0.0952 when χ = 0.1 and *n* = 1000. This means that by collapsing branches shorter than 0.1/ 1000 in the Bayesian bootstrap trees, the theoretical support of branches corresponding to 1 mutation in the reference tree will be 0.9048. In short, collapsing branches shorter than χ(= 10%)/ *n* gives support very close to 1−χ(= 90%).

Now, let us examine how this analysis works in the more general case where a branch of the reference tree corresponds to *k* mutations. In this case, we consider the sum *S*_*k*_ of *k* weights, each distributed according to the same law (see above), and we search for the probability that *S*_*k*_ is less than χ and that the corresponding branch is collapsed. Using the exponential approximation (fully justified in practice, since *n* is rarely less than 100), we are interested in the distribution of a sum of exponential laws with parameter 1.0, which is an Erlang law whose distribution function is easy to compute. We find theoretical supports of 0.9048, 0.9953, 0.9998, and 0.9999 for *k* = 1, 2, 3, and 4 mutations.

We thus have a method to approximate Bayesian supports as a function of branch length. We will see on large simulated and real data sets that this approximation is close to the values of Bayesian support observed for correct branches, when the data do not deviate drastically from the perfect phylogeny assumptions. Compared to the standard frequentist framework, an advantage of the Bayesian approach is that we can control this support to make it meaningful and interpretable in practice by adjusting the collapse threshold. The simulation results below confirm the relevance of the approach, which produces supports close to standard confidence levels with low homoplasy data.

### Criteria for evaluating and comparing branch supports

In the following, we evaluate and compare different branch supports on simulated and real data.

We describe here the main principles and criteria used in these analyses.

A basic criterion is that a support is relevant if it allows a certain separation between correct branches (with higher support) and false branches (with lower support). Of course, this criterion is not sufficient. For example, branch length is correlated with branch correctness, but length is not a support in the usual sense. One reason (but not the only one) is that branch lengths are neither normalized (between 0.0 and 1.0) nor interpretable. For example, with low homoplasy data, branches representing very few mutations are generally correct (Wertheim et al. 2022), although their length (number of mutations divided by sequence length) is very close to 0.0. For it to be interpretable, we want a support to be close to a confidence level of a standard statistical test, where strong supports are close to 1.0.

In statistical testing, we distinguish between type 1 error rate (i.e., the probability of selecting an incorrect branch) and type 2 error rate (i.e., the probability of not selecting a correct branch). Power is often used, which is equal to 1.0 minus the type 2 error rate (i.e., the probability of selecting a correct branch). Type 1 error rate and power are in tension: with a high selection threshold, there will be few type 1 errors, but power will be low, and vice versa. A classic measure of the performance of a support (or test statistic or discriminant function) is the AUC under the ROC curve (Area Under the “Receiver Operating Characteristic” Curve), which combines the type 1 error rate and power and integrates results over all possible selection thresholds between 0% and 100% and is not dependent on any particular threshold. We use this criterion with simulated data to quantify the performance of different bootstrap supports (see Ecker et al. 2024, for the use of ROC-AUC with branch supports).

In terms of interpretability, to come close to a usual confidence level, we aim to guarantee that when the support has a value greater than or equal to (typically) 95%, the type 1 error rate is at most 5%. In our experiments with simulated data, we set the type 1 error rate to 5% (or 10%) and check that the corresponding selection threshold is equal to or less than 95% (or 90%) but close to this value to avoid the support being overly conservative (i.e., rejecting most branches and selecting few correct ones). To quantify this effect, we calculate the power associated with a given risk level (5% or 10%), which should be as high as possible (see Berry and Gascuel 1997, and Anisimova et al. 2006, 2011, for the use of these measures with branch supports).

In the following, these criteria are evaluated on simulated data that gradually deviate from the perfect phylogeny. These data are used to test the robustness of our theoretical predictions based on the perfect phylogeny. If these predictions are robust, the expected value of the support for a correct branch (e.g., ∼0.632 or higher, for the frequentist bootstrap with collapse) gives an indication of the selection threshold to be adopted to separate correct branches from false branches with lower support.

Real data is different from simulated data. In particular, we usually do not know which branches are correct and which are false, and the above criteria are usually not measurable. But with viruses (and other data sets) we are in principle relatively close to perfect phylogeny. We therefore expect real data with low homoplasy to resemble our simulated data, and results obtained on simulated data to be largely transposable to real data. In the following, we use several metrics and criteria to measure this proximity: (i) The distribution of branch lengths in the inferred tree; this length is expressed in number of mutations (equal to the estimated branch length multiplied by the MSA length), with peaks expected around integer values of the number of mutations, which should be low (Fig. 2). (ii) The level of homoplasy in the MSA, where we compare the total length (in number of mutations) of the tree, to the minimum number of mutations required to explain each of the sites in the MSA (e.g., a site with 2 different nucleotides requires at least 1 mutation; see below for details and examples). (iii) The robustness of our theoretical support for branches as a function of their lengths (in number of mutations; Fig. 3). We will see that the analyzed datasets (EBOV, SARS-CoV-2…) are close to the simulated datasets according to criteria (i) and (ii), and that (iii) the obtained supports are similar to the theoretical predictions, thus supporting the application to real data with moderate homoplasy of the selection thresholds obtained by theoretical analysis and simulation.

**Figure 3:**
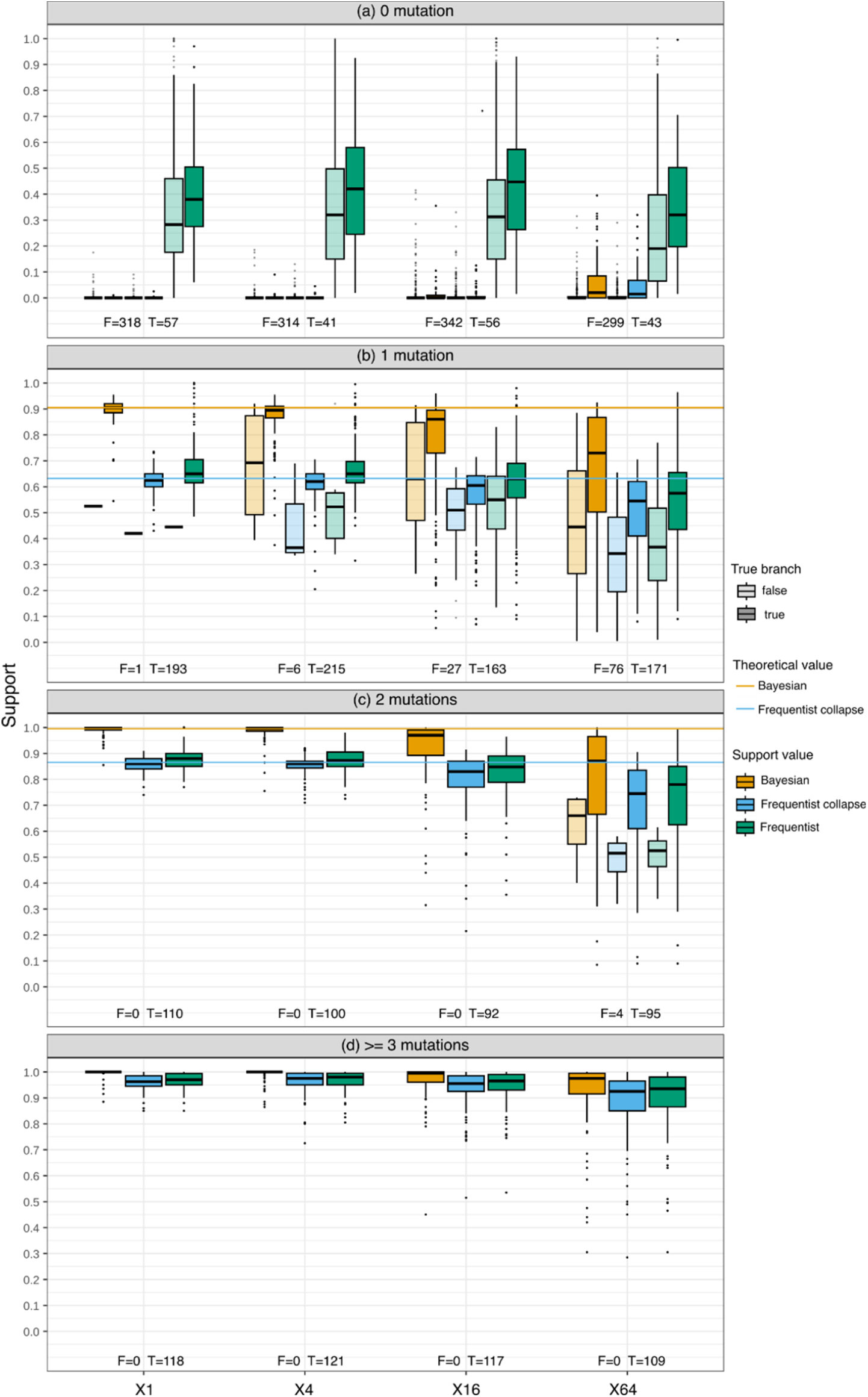
Simulation results. X-axis: the four experimental conditions: X1 (EBOV-like, low homoplasy: 14%), X4 (moderate homoplasy: 60%), X16 (substantial homoplasy: 222%), and X64 (high homoplasy: 518%). Y-axis: branch supports. Panels **(a)** to **(d)**: supports of branches with estimated lengths in the range [0, 0.5[, [0.5, 1.5[, [1.5, 2.5 [and [2.5, ∞[, respectively. Frequentist: standard frequentist bootstrap, where near-zero bootstrap branches are not collapsed. FrequentistCollapse: frequentist bootstrap, where bootstrap branches with length corresponding to <0.1 mutation are collapsed. Bayesian: our approach with weighted sites, where bootstrap branches with length corresponding to <0.1 mutation are collapsed. Within each panel, we provide the number of false (F) and true (T) branches, for each experimental condition (e.g., F = 299 and T = 43 for 0 mutation and X64 in panel (a)). We also provide the theoretical support values for the FrequentistCollapse and Bayesian approaches, under perfect phylogeny assumptions. See text for details.

## Results

### Simulated data

To evaluate the properties of the various approaches and the robustness of our theoretical results with large realistic datasets, we simulated sequences inspired by the Ebola virus (EBOV) dataset (Dudas et al. 2017; Zhukova et al. 2023) that is analyzed below (see also Fig. 1). The complete EBOV MSA consists of 1,610 sequences and 18,996 sites. For computational purposes, we randomly subsampled this dataset to select 800 sequences. We inferred a tree from these 800 sequences using PhyML+GTR+G4 (Guindon et al. 2010). This “EBOV” tree has a total length of 0.0926, corresponding to ∼1760 mutations (i.e. ∼1.10 mutation per branch). We used the EBOV tree to simulate the evolution of sequences. However, to avoid having too many zero-length branches with simulated data, we replaced all near-zero branch lengths in the EBOV tree with values drawn from an exponential distribution with an expectation corresponding to 1/2 mutation. Such correction is required because in simulations short branches can generate zero mutation, even if their length is larger than zero. The resulting tree (same topology, slightly longer length) is called “CorrectedEBOV” tree in the following.

We rooted the CorrectedEBOV tree using mid-point rooting and took as ancestral sequence the sequence made up of the most common nucleotide at each site in the original alignment. This root sequence was evolved along the tree branches using the GTR+G4 model estimated by PhyML with EBOV sequences. We then inferred a tree from the simulated sequences using PhyML+GTR+G4. This “Simulated” tree is very close to the EBOV tree. When collapsing the branches with near-zero length, the Simulated tree has 421 internal branches in common with the CorrectedEBOV tree, and only 1 specific internal branch. This confirms that short trees are not fully resolved (with many zero-length branches), but almost all non-zero-length branches are correct (Fig. 3). We also measured the level of homoplasy in this dataset. Based on the simulated MSA, the minimum number of mutations is equal to 1,942. To obtain this result, we assume a perfect phylogeny and count for each site the number of nucleotides present in the site and subtract 1 (e.g., 3 nucleotides = 2 mutations); then we sum the values thus obtained for all sites. The actual size (in number of mutations = sum of branch lengths x alignment length) of the tree inferred from the simulated MSA is 2,217, which (as expected) is larger than the theoretical minimum, due to homoplasy, convergence, reversion, etc. However, both values are close with 14% additional mutations (= (2217 −1942) 1942)). The same measure applied to the EBOV MSA and tree is equal to 15%, indicating the closeness of the two datasets in terms of homoplasy. Finally, the branch lengths in the Simulated tree show a similar distribution to that of the EBOV tree, with very clear peaks corresponding to 0, 1, 2, 3 … mutations (Fig. 2). However, as expected, the peaks are slightly less pronounced in the real data than in the simulated data, especially when the number of mutations is large, with a few intermediate lengths corresponding to non-integer numbers of mutations.

To study the properties and limitations of the approaches, we simulated MSAs and trees that gradually deviate from the assumptions of perfect phylogeny up to conditions where these assumptions are clearly violated. To achieve this we multiplied the branch lengths in the CorrectedEBOV tree by a constant factor, and divided the number of sites in the MSA by the same factor. In this way, the number of expected mutations remains constant, i.e. the phylogenetic signal remains similar, but the degree of homoplasy increases and one gets further and further away from a perfect phylogeny. The factors used in our simulations were equal to 4, 16 and 64. For example, with a factor of 64, the tree length is close to 6.0 mutations per site (instead of 0.0926), while the length of the MSA is 296 (instead of 18,996), a radically different setting corresponding to, for example, animal or plant genes rather than virus genomes. Sequence simulation and tree inference were then performed using the same programs and options as for the original CorrectedEBOV tree. The estimated level of homoplasy was equal to 60%, 222%, and 518% for factors of 4, 16, and 64, respectively, a clear violation of the assumptions of perfect phylogeny (= 0%). In the following, the settings corresponding to the different multiplicative factors are called “X1” (i.e. the original setting corresponding to EBOV data), “X4”, “X16” and “X64”.

Using PhyML+GTR+G4, we applied three bootstrap methods to each of these simulated MSAs and trees: (1) The original frequentist approach of Felsenstein (1985), where all branches in the bootstrap trees are kept, even if they have nearly zero length (called “Frequentist” in the following). (2) The frequentist approach where bootstrap branches of length < 0.1 /*n* are collapsed (called “FrequentistCollapse” in the following; *n* is the length of the MSA, see explanations above). (3) Our Bayesian approach where the sites are weighted and the bootstrap branches with length < 0.1/ *n* are collapsed (called “Bayesian” in the following). For each of the four simulated trees (corresponding to X1, X4, X16, and X64) and each branch of these trees, we thus obtained three supports corresponding to the three bootstrap methods. The branches were classified depending on their length in the Simulated trees. The “0 mutation” branches have length estimates less than 0.5/ *n*, the “1 mutation” branches have length estimates in the range [0.5, 1.5 [/*n*, the “2 mutations” branches in the range [1.5, 2.5[/ *n*, etc. Indeed, we do not expect the estimated branch lengths to correspond exactly to 0, 1, 2 … mutations, although the distribution of branch length estimates (Fig. 2) shows that we are relatively close to this pattern, especially with the EBOV-like (X1) data and tree. In addition, we distinguished the “true” and “false” branches in trees inferred from simulated data, depending on whether the corresponding bipartitions are found in the model (EBOV) tree or not. The results are shown in Figure 2 and 3.

As we approach perfect phylogeny, the estimated branch lengths (multiplied by the MSA length) are remarkably close to an integer number of mutations (Fig. 2(a), X1). This trend can also be seen in the analysis of real virus and low homoplasy data (Fig. 2(c-e); see below). As expected, this trend diminishes as we move away from perfect phylogeny and the phylogenetic signal is perturbed (Fig. 2(b), X64). Under these conditions, the peaks of the distribution over integer values remain visible, but they are less pronounced due to the high level of homoplasy.

As expected, when the estimated length corresponds to ∼0 mutations (Fig. 3(a)), the inferred branches are generally false (∼85% or more), justifying the low support of the FrequentistCollapse and Bayesian bootstraps, even if 12% to 15% of the inferred branches (mostly corresponding to cherries) are true by chance. These approaches indicate that there is no signal in the data for these branches. We could think about alternatives, such as using some prior on the resolution of multifurcations (e.g., giving a support of 1/3 in the case of trifurcation), but this is beyond the scope of this article, where we consider that the goal of branch supports is to measure the signal in the data. The standard Frequentist bootstrap provides larger supports (sometimes close to 100%) for both true and false branches that are indistinguishable by their support values, although the median support of the true branches is slightly higher than that of the false branches. For example, using the standard support threshold of 70%, no branches are supported by Bayesian and FrequentistCollapse, but 29 (vs 5), 33 (vs 6), 28 (vs 9), and 26 (vs 3) false (vs true) branches are retained by Frequentist in the X1, X4, X16, and X64 conditions, respectively. Thus, with a threshold of 70%, the Frequentist bootstrap supports a large majority (∼75% in total) of false branches. This confirms (see also Fig. 1) the importance of collapsing near-zero branches and using multifurcating trees, regardless of the type of bootstrap support.

With an estimated length of ∼1 mutation (Fig. 3(b)), the signal becomes worse (as expected) when the multiplicative factors are high: with X1 we find 1 false branch out of 194 (∼0.5%), while with X64 ∼31% of branches are false. The Bayesian support is significantly higher than the Frequentist support, which is slightly higher than FrequentistCollapse (as expected, see above). For X1, X4, and to some extent X16, the median (FrequentistCollapse and Bayesian) supports of the true branches are close to the theoretical values (∼63% and ∼90%, respectively), although we are relatively far from a perfect phylogeny, especially for X16 (homoplasy = 220%). For X64 (very far from perfect phylogeny, homoplasy = 518%), we deviate from the theoretical values, and the observed supports of the true branches are significantly lower than the predictions. In all conditions and with the three bootstrap versions, the supports of the false branches are significantly lower than those of the true branches, justifying (if necessary) the use of the bootstrap. The length of branches gives a good indication of their support, but cannot be used alone, for example in the presence of contradictions between sites.

With an estimated length of ∼2 mutations (Fig. 3(c)), very few branches are false (4 in total for X64). The difference in support between the two Frequentist versions and the Bayesian bootstrap is still significant, and Frequentist is again slightly higher than FrequentistCollapse. The values observed for X1, X4, and to some extent X16 are still close to the theoretical values (∼86% and ∼99%, respectively) derived from perfect phylogeny assumptions. Again, the support values for X64 deviate significantly from the theoretical values, and for the three approaches, the 4 false branches have much lower bootstrap support than the true branches.

With an estimated length of ∼3 mutations and more, there are no longer any false branches, and the supports of the three approaches come very close to 1.0, even for X64.

To summarize the observations in Figure 3: (1) It is essential to collapse null branches and consider multifurcating trees (see also Fig. 1). (2) With low homoplasy, the Bayesian support is higher and more interpretable than the Frequentist (collapsed or not) support, with values close to 100% for true branches. (3) The supports are similar and close to 100% for long branches, which are generally true. Moreover, as expected, the Bayesian and Frequentist (collapsed or not) supports yield substantially different values but are strongly correlated for non-zero length branches (Pearson correlation in the range [80%-94%] in the 4 simulated conditions for Bayesian vs Frequentist and Bayesian vs FrequentistCollapse).

We compared the performance of the three approaches using ROC-AUC. We grouped the results for all four conditions to obtain an overall view, and because the number of false positives for X1 and X4 is very low. We also collapsed the unresolved parts of the inferred reference trees for all the reasons mentioned above (Fig. 1 and Fig. 3; only the results in panels (b), (c), (d) of Fig. 3 were retained). The ROC curves in Figure 4 show a slight but significant advantage for the Bayesian bootstrap (ROC-AUC = 0.918) over the two Frequentist versions (0.895 and 0.903, for standard and collapse, respectively).

**Figure 4:**
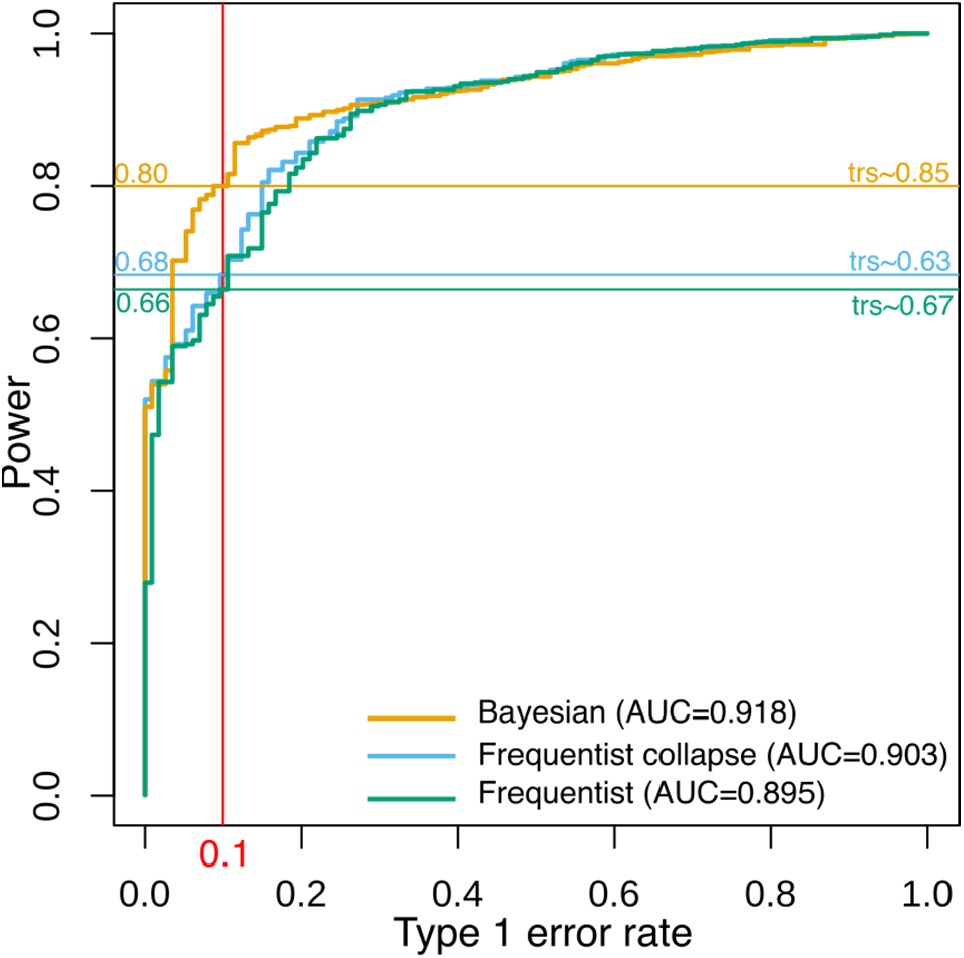
ROC curves and AUC of the three bootstrap approaches on simulated data. The four simulation conditions (X1 to X64) are grouped together and we use the multifurcating version of the inferred reference trees, where near-zero branches are collapsed. With a fixed level of risk of 0.1 (10%), we show the power (e.g., 0.80 for the Bayesian bootstrap) and the selection threshold (e.g., trs=0.85).

In practice, bootstrap users want to know the threshold at which branches should be considered well supported. In the case of the standard Frequentist bootstrap, an empirical threshold of 70% is commonly used, as suggested by Hillis and Bull (1993), Soltis and Soltis (2003), and others. For the Bayesian bootstrap, our theoretical analyses above suggest a threshold of ∼90%, which is easier to interpret because it is analogous to the confidence level of a statistical test. The results in Figure 4 show the threshold and power for the three approaches and all four conditions grouped together (with low to high homoplasy), when the risk level is fixed at 10%. The two Frequentist versions have similar power (66-68%), with a higher selection threshold (as expected) for the standard version (67%) than for the collapse version (63%, similar to theoretical prediction, despite high homoplasy of some datasets, especially X64). The Bayesian bootstrap obtains a significantly higher power (80%; p-value ≈ 0.0 using Mac Nemar test) with a threshold relatively close to the theoretical prediction (85% instead of 90%, explained by the clear departure from perfect phylogeny with X64; Fig. 3). When the risk level is fixed at 5%, we still observe a significant gain in power (59% and 70%, respectively, for the Frequentist and Bayesian approaches; p-value ≈ 0.0), while the selection thresholds become close to 70% for the Frequentist versions (standard: 72%, collapse: 65.5%) and close to 90% for the Bayesian bootstrap (88.5%). Once again, we see the superiority of the Bayesian approach, with better interpretability and higher power for the same level of risk.

To summarize the results of these experiments: (1) Across the four conditions (with very different levels of homoplasy), the Bayesian bootstrap performs better in terms of ROC-AUC and power. (2) We recover the usual 70% threshold for the standard Frequentist bootstrap, although our simulation protocol is radically different from that used in (Hillis and Bull 1993), with small model trees of 4 to 9 taxa, whereas we had 800. (3) The collapse version of the Frequentist bootstrap performs slightly better than the standard version in terms of ROC-AUC, with an optimal selection threshold of ∼65%, close to the theoretical prediction. (4) A 90% threshold for the Bayesian bootstrap is reasonable, with a low type 1 error rate (∼3.5%), but a lower threshold of 85% will provide more power, depending on the level of risk accepted. Note that all these results were obtained using the multifurcating version of the inferred reference tree, in which near-zero branches inferred by chance were collapsed (and not counted in the results). The two Frequentist versions differ in whether or not they collapse the bootstrap trees, which has a moderate effect when the reference tree is already collapsed. In fact, IQTREE’s ‘czb’ option collapses both the reference and bootstrap trees, while RAXML-NG’s ‘bestTreeCollapsed’ collapses only the reference tree.

### Results with virus and low homoplasy datasets

We used three virus datasets, from Ebola in West Africa (Dudas et al. 2017), from the global SARS-CoV-2 pandemic (Zhukova et al. 2021), and from Rift Valley fever virus (RVFV) downloaded from NCBI Genbank. We also used one large low-homoplasy plant MSA collected from TreeBASE (Piel et al. 2009; Togkousidis et al. 2023). Details on all these datasets (and the simulated ones) are provided in Supp. Table 1.

The West Africa Ebola epidemic began in 2013 in Guinea and circulated for two and a half years in neighboring Liberia and Sierra Leone. We downloaded the Ebola virus (EBOV) full-genome alignment of (Dudas et al. 2017) from (https://github.com/evolbioinfo/bdei). This multiple sequence alignment (MSA) consists of 1,610 sequences of 18,996 nucleotides. From this MSA, we randomly subsampled 800 sequences for computational reasons. We inferred the reference tree using PhyML with GTR+G4 and default options. To infer frequentist bootstrap trees, we generated 200 bootstrap alignments by sampling sites with replacement (as usual) and used PhyML+GTR+G4. To infer Bayesian bootstrap trees, we generated 200 weight vectors using a Dirichlet distribution with parameters (18996; 1,…,1) and used PhyML, which allows for continuous site weights, with GTR+G4 and default options. All these steps were performed using our BBOOT pipeline (see Supp. Mat. for details). The Frequentist supports were computed as usual, while for the FrequentistCollapse and Bayesian supports, we first collapsed the bootstrap branches with length less than 0.1 (mutation) 18996 (MSA length). In all this, we used the very same approach and program options as for the simulated data, thus making the results comparable to the X1 (EBOV-like) condition. In fact, the amount of homoplasy is similar for the two datasets (∼15%).

Our SARS-CoV-2 dataset covers the first five months of the pandemics (up to April 25, 2020). It was subsampled to avoid over-representation of certain countries and periods (Zhukova et al. 2021). We extracted the identifiers from the publication and downloaded the sequences from GISAID. We again randomly subsampled the sequences for computational reasons and aligned them to obtain an MSA of 796 sequences with a length of 29,903, representing the emergence of the global pandemic. Alignment sites known to contain noisy phylogenetic signal were masked. The reference and bootstrap trees were inferred using the same protocol as for the EBOV MSA, with a collapse threshold of 0.1 29903. The size of the tree (in number of mutations) is 1062, while assuming a perfect phylogeny, the lower bound for the number of mutations is 787, resulting in a homoplasy level of 35%. This is appreciably higher than for the EBOV dataset (15%), although the number of mutations is significantly lower (∼0.67 mutations per branch versus ∼1.10 with EBOV).

Our RVFV dataset was downloaded from Genbank (on April 17, 2024) and consists of 300, 240, and 237 sequences of its three respective segments (S, M, and L). We aligned the three segments separately to obtain three MSAs of length 1690, 3885, and 6404 nucleotides for the S, M, and L segments, respectively. The reference and bootstrap trees were inferred using the same protocol as above, with a collapse threshold of 0.1 /MSA length. The lengths of the three trees in number of mutations are 955, 2177, and 2976 for S, M, and L segments (i.e. ∼1.6, ∼4.6 and ∼6.3 mutations per branch), while assuming a perfect phylogeny, the lower bounds for the number of mutations are 505, 1302, and 1778, resulting in estimated homoplasy levels of 89%, 67%, and 67%, respectively. This is significantly higher than for the two previous data sets (15% and 35%) and similar to the simulated X4 condition. The branch lengths and supports of the three trees were combined in the results.

To investigate the behavior of bootstrap supports with non-viral data, we used a large low-homoplasy dataset collected from TreeBASE (Piel et al. 2009; Togkousidis et al. 2023; Supp. Mat.). We used the largest nucleotidic MSA with a homoplasy level of less than 50%, containing 385 (268 unique) sequences from the genus *Psiguria* (*Cucubirtacea*, flowering plants). This genus has 6 species, the dataset is both inter and intra specific. The number of sites is 1140 (much less than EBOV and SARS-CoV-2, but similar to RVFV segment S). The number of mutations per branch is 1.1 (similar to viruses) and the level of homoplasy is 46%, which is relatively high compared to viruses. The reference and bootstrap trees were inferred using the same protocol as for viruses.

The results for these four datasets are displayed in Figure 2 (distribution of branch lengths) and Figure 5 (branch supports). We observe:

**Figure 5:**
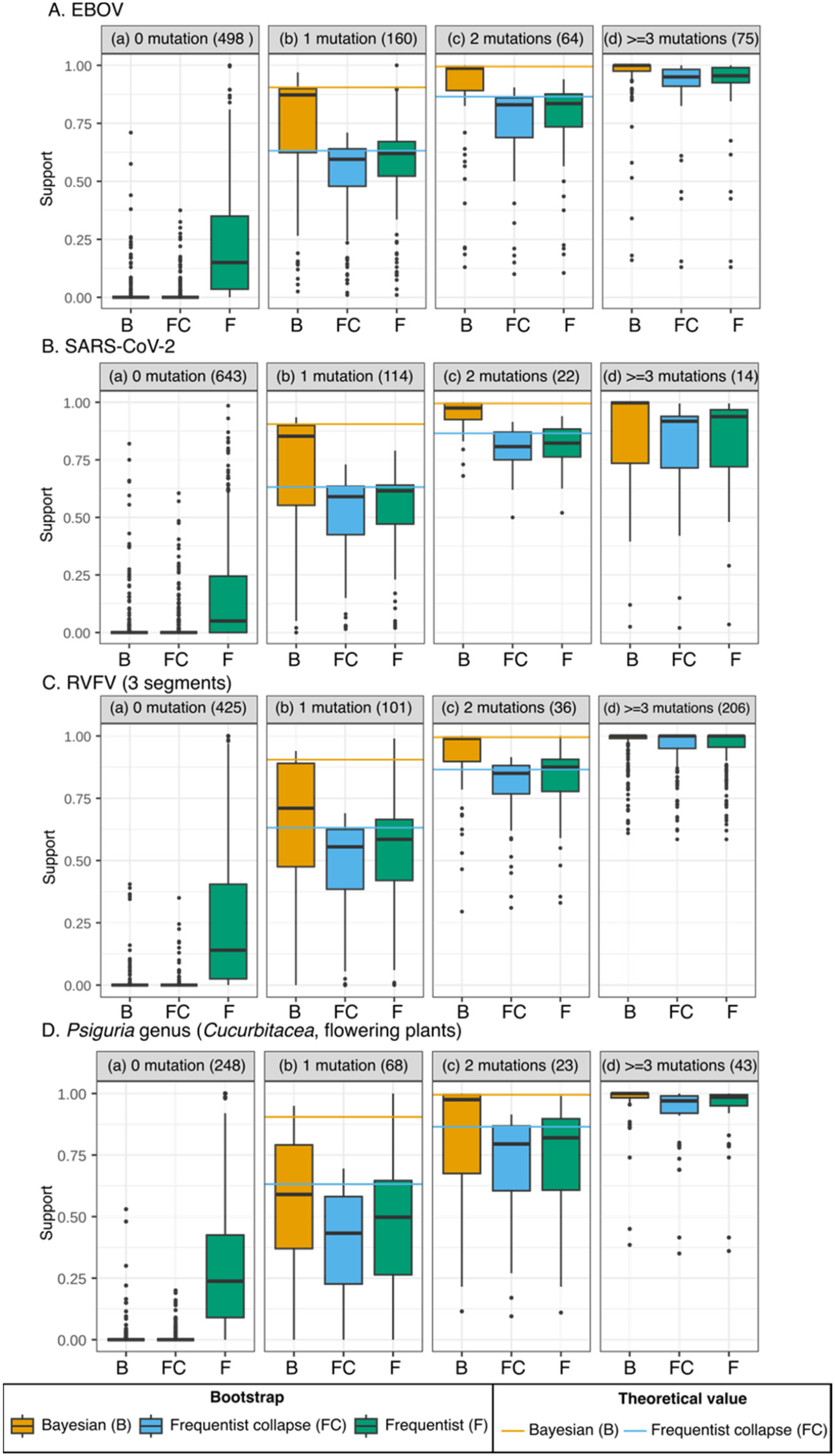
Analysis of viral and non-viral datasets. **A**: EBOV (Dudas et al. 2017); **B**: SARS-CoV-2 (Zhukova et al. 2021); **C**: Rift Valley fever virus (RVFV; extracted from Genbank); **D**: *Psiguria* genus (plants) low-homoplasy dataset collected from TreeBASE. Panels **(a)** to **(d)**: supports of branches with estimated lengths in the range [0, 0.5[, [0.5, 1.5[, [1.5, 2.5[, and [2.5, ∞[, respectively. Frequentist: standard frequentist bootstrap, where short bootstrap branches are not collapsed. FrequentistCollapse: frequentist bootstrap, where bootstrap branches with length corresponding to <0.1 mutation are collapsed. Bayesian: our approach with weighted sites, where bootstrap branches with length corresponding to <0.1 mutation are collapsed. Within each panel, we provide the theoretical support values for the FrequentistCollapse and Bayesian approaches, under perfect phylogeny assumptions. See text and Supp. Table 1 for details.

The branch lengths are similar between the simulated data in condition X1 and the real EBOV data (Figures 2(a) and 2(c)), with marked peaks on the first integer values corresponding to 0, 1, 2, and 3 mutations. However, with the real data (Fig. 2(c)), there are intermediate points between these integer numbers of mutations, and the graph becomes more blurred beyond 3 mutations. This is a sign of a less perfect phylogenetic signal than with the simulated data, in particular due to some violation of the substitution model (GTR+G4) used to infer the tree and estimate branch lengths. With SARS-CoV-2, there are fewer mutations than with EBOV, and many branches of zero length (∼80%). There are peaks at 0, 1 and 2 mutations, but a substantial number of intermediate points, particularly between 0 and 1 mutations. This indicates an increased deviation from the perfect phylogeny, as indicated by the homoplasy level (35%), due to misspecification of the substitution model and probably also because convergences in these sequences were observed very early (Van Dorp et al. 2020), thus perturbing the phylogenetic signal. Despite a rather high homoplasy level (>65%) in the RVFV datasets, we still see relatively well-marked peaks for 0, 1, 2, and 3 mutations, but the values are less well centered on integer values than for EBOV. The same can be seen for the plant dataset (homoplasy = 46%), the peaks are visible with 0, 1, 2, and 3 mutations, but they tend to be shifted to the right due to model violation and the presence of duplicated sequences, gaps and unknowns that blur the signal and make it difficult to assign a mutation to a unique branch in the tree.

Despite these differences in branch length distribution and a noticeable deviation from perfect phylogeny, the branch supports (Fig. 5) are close to the theoretical values and to the results obtained with simulated data (Fig. 3). We see again (Fig. 5A-B-C-D(a), ∼0 mutation) the importance of collapsing the short branches of the inferred trees, without which many branches not supported by any mutation have substantial support using the standard Frequentist bootstrap. For some datasets (e.g. SARS-CoV-2, Fig. 5B(a)), there are still near-zero branches that receive FrequentistCollapse and Bayesian support quite far from 0.0, presumably because of their intermediate length, but these supports remain substantially below the selection thresholds of 63-65% (FrequentistCollapse) and 85-90% (Bayesian) obtained with simulated data; most of these branches are probably incorrect.

For branches corresponding to ∼1 mutation, the median supports for both the EBOV and SARS-CoV-2 datasets are very close to the theoretical values. Again, the standard Frequentist bootstrap has slightly higher supports than the collapse version. By using thresholds of 85% for the Bayesian bootstrap and 63-65% for the Frequentist versions, we are very likely selecting branches that correspond to the true evolutionary history of these viruses. For RVFV and plant trees, branches corresponding to ∼1 mutation have FrequentistCollapse and Bayesian supports below the theoretical values (Fig. 5C-D(b)), as expected given the relatively high level of homoplasy in these data. With thresholds of 85% (Bayesian), 65% (Frequentist), and 63% (FrequentistCollapse), very few branches of length ∼1 are selected, which is likely conservative despite the rather low level of these thresholds.

For branch lengths of ∼2 mutations and more, the results are consistent with the theoretical and simulation results for all data sets (Fig. 5A-B-C-D(c-d)), and the use of thresholds of 90% (Bayesian), 70% (Frequentist) and 65% (FrequentistCollapse) should ensure the selection of mostly correct branches. However, it is not possible to know (as with the simulated data) whether one support gives better accuracy than the other. It is likely that their performances are relatively close (their Pearson correlation is in the range [88%-94%] on the 4 datasets), but the Bayesian bootstrap shows again the advantage over the Frequentist versions of corresponding to thresholds that are more interpretable in the usual sense of statistical testing.

## Discussion

The Bayesian bootstrap is a smoothed version of the frequentist bootstrap in which the sampling with replacement procedure is replaced by weights on the statistical units (or observations) drawn from a non-informative distribution (Rubin 1981; Lo 1991; Shao and Tu 1995). Here, we have investigated the application of this principle to the estimation of the branch supports of a phylogeny. Our results demonstrate the interest of this smoothed approach for short trees and branches inferred from highly conserved data sets with low numbers of mutations, as we observe in viruses (but not only). Such datasets contain a partial but reliable evolutionary signal, with parts of the inferred phylogenies that are poorly resolved, but few false branches (Wertheim et al. 2022). Our theoretical and experimental results on simulated and real data show that Bayesian supports are higher than standard frequentist supports, and more consistent with the usual statistical interpretation of a test. A selection threshold of 85-90% gives good results on sequences with low homoplasy, with very few false positives and fairly good power. To achieve similar performance with the standard bootstrap, a selection threshold of 65-70% must be used, depending on whether near-zero branches in the bootstrap trees are collapsed or not. In our simulations, the Bayesian bootstrap shows better accuracy (ROC-AUC, power for a given risk level, Fig. 4), probably due to its smoothness. Our results also show that the two supports are strongly correlated, and that as the trees and branches lengthen, with more mutations and a high level of homoplasy, the two supports become very close (as expected from bootstrap theory; Efron and Tibshirani 1993).

For this type of data with low homoplasy, bootstrap supports are strongly correlated with branch lengths (Felsenstein 1985). However, the usefulness of the bootstrap and statistical branch supports in general remains, especially for short branches corresponding to 1 or 2 mutations, where the signal can be conflicting and blurred by the homoplasy of the data, even if it is small (Fig. 3(b), 3(c)). Our results show that to properly interpret the results of a bootstrap analysis, it is important to measure the level of homoplasy in the data and to plot the distribution of branch lengths to verify that the lengths cluster (or do not cluster) around integer values of the mutation number (Fig. 2). These statistics (homoplasy level, branch length distribution) and the branch supports of the Bayesian bootstrap can be obtained using our pipeline BBOOT (https://github.com/evolbioinfo/bayesian_bootstrap) based on PhyML for tree inference (Guindon et al. 2010). The computing time for the Bayesian phylogenetic bootstrap is the same as for the standard approach.

To compare the behavior of frequentist and Bayesian bootstrap supports on a wide variety of sequences, we applied these approaches to large datasets collected from TreeBASE (Piel et al. 2009; Togkousidis et al. 2023; Supp. Mat.). The results for DNA (300 datasets, Supp. Fig. S1) show that: (1) homoplasy levels in these datasets are highly variable, from less than 1% to over 30,000%, with an average of 1,210%; we are therefore often far from the homoplasy levels studied here (between 15% and 89%; Supp. Table 1), but these datasets are peculiar, virus-free (except for one Dengue MSA) and often from deep phylogenies. (2) Branch support differences between the two bootstrap approaches depend on the level of homoplasy (as expected). The differences are small (<10%) when homoplasy exceeds 1000%, but Bayesian supports remain higher than frequentist supports. (3) For each of the DNA data sets, the correlation between the two branch supports is very high (≥93%, 98% on average). The results are similar for proteins (100 datasets, Supp. Fig. S2), but we did not find any large dataset with low homoplasy (minimum value = 182%, average = 1,351%), probably because protein datasets in TreeBASE were generally used to elucidate the deep branches of the tree of life. Again, we observe that the gap between the two supports narrows with high homoplasy (<6% with >1000% homoplasy), and that the correlation between the two supports is strong for all datasets (>95%, 99% on average).

Taken together, our results show that the standard frequentist version of the bootstrap would benefit from being replaced by the Bayesian version, at least in the case of DNA data. The Bayesian bootstrap is smoother, its supports are higher and therefore less conservative, especially when there are few mutations and homoplasy is low. Several avenues are open for future research. In particular, our approach is Bayesian in that it assumes an a priori distribution on the observations (the MSA sites), but it differs markedly from the standard Bayesian approach in phylogenetics, where we usually have an a priori distribution on the parameters to be estimated, but not on the sites. The standard Bayesian approach is based on the likelihood and posterior probability of the inferred trees, but does not question the data provided as input, as we do here by weighting the sites. The standard Bayesian supports are high and generally liberal, whereas the standard bootstrap gives generally conservative low supports (Douady et al. 2003). It would be interesting to compare the performance of our Bayesian supports with the usual Bayesian posteriors, in particular aBayes, which is fast and fairly accurate (Wertheim et al. 2022; Ecker et al. 2024), and possibly to combine the two approaches to obtain better performing and easily interpretable supports. As mentioned above, our Bayesian approach, designed to reveal the signal in the data, could be complemented by priors on tree topologies and shapes (e.g., coalescent, birth-death). Another issue with the phylogenetic bootstrap is the computing time; fast algorithms for the Bayesian version would be very useful, in continuation of the Rapid and Ultrafast bootstrap algorithms developed for the frequentist version (Stamatakis et al. 2008; Minh et al. 2013). Lastly, the Bayesian bootstrap could be applied to other types of supports and statistics in phylogenetics, in particular the Transfer Bootstrap Expectation (TBE; Lemoine et al. 2018), which measures the degree of presence of reference branches in bootstrap trees, and the RELL bootstrap, which is used to compare and test phylogenetic trees (Kishino et al. 1990; Goldman et al. 2000; Shimodaira and Hasegawa 2001).

## Data availability

All our datasets and workflows are available from (see Supp. Mat. for details): https://github.com/evolbioinfo/bayesian_bootstrap_analyses/.

## Acknowledgements

The authors would like to thank Stéphane Guindon for his help with PhyML and Marie Morel for early discussions and preliminary analyses.

## Supplementary Material

## Material and methods

### Theoretical dataset

- EBOV multiple sequence alignment (800 sequences from the EBOV dataset below) was deduplicated using goalign dev0537492, and R v4.3.3:
  - Ambiguous characters were replaced by gaps:

~~~
goalign replace -s ‘N’ -n ‘-’ -p -i <MSA> | goalign replace -s ‘?’
-n ‘-’ -p | goalign replace -s ‘M’ -n ‘-’ -p | goalign replace -s ‘W’ -n ‘-’ -p | goalign replace -s ‘R’ -n ‘-’ -p | goalign replace -s ‘K’ -n ‘-’ -p | goalign replace -s ‘Y’ -n ‘-’ -p > msa_ambig
~~~
  - Distance matrix was computed without considering difference between gaps and characters: goalign compute distance --gap-mut 0 -i msa_ambig -p -t 10 -m pdist > dist
  - The distance matrix was used to extract one sequence per identical group of sequences using connected components computed with “concom” R package.
- Sequences were then selected by names and gap-only sites were removed using goalign dev0537492

~~~
goalign subset -p -i <MSA> -f <NAMES> | goalign clean sites -c 1 – p > <OUT MSA>
~~~
- Phylogenetic tree with frequentist (collapse) bootstrap supports was inferred using IQTREE v2.2.2.5

~~~
qtree -s <MSA> -m GTR+G4 --seed 123456789 -b 200 (-czb)
~~~
- PhyML Bootstrap alignments were computed using goalign dev0537492

~~~
goalign build seqboot -S -p -n 200 -i <MSA> -o <PREFIX>_ --seed 123456789
~~~
- RAxML Bootstrap alignments were computed using goalign dev0537492

~~~
goalign build seqboot -S -p -n 200 -i <MSA> -o <PREFIX>_ --seed 987654321
~~~
- Phylogenetic trees (reference + frequentist bootstrap) for the deduplicated EBOV multiple sequence alignment were inferred using RAxML-NG v1.1.0:

~~~
raxml-ng --msa <MSA> --model GTR+G4 –seed 123456 --threads 1
~~~
- Phylogenetic trees (reference + frequentist bootstrap) for the deduplicated EBOV multiple sequence alignment were inferred using PhyML v3.3.20200621:

~~~
phyml -i <MSA> -m GTR -c 4 -a e -f e -d nt -o tlr -b 0 --r_seed 123456
~~~
- Frequentist bootstrap supports from non-collapsed bootstrap trees were computed using gotree devb324e73:

~~~
gotree compute support fbp -i  -b <BOOT> -o <OUT TREE WITH SUPPORTS>
~~~
- Bayesian and frequentist Bootstrap supports from collapsed bootstrap trees were computed using gotree devb324e73:

~~~
gotree collapse length -i <BOOT> -l <0.1/length> | gotree compute support fbp -i  -b --o <OUT TREE WITH SUPPORTS>
~~~

### EBOV dataset

- Ebola virus dataset was downloaded from Github:
  - repository : https://github.com/evolbioinfo/bdei
  - file : bdei/ebola/data/aln.ids.fa
- 800 sequences from this MSA were randomly sampled using goalign dev0537492:

~~~
goalign sample seqs -n 800 -i ${msa} –p.
~~~
- Phylogenetic trees (reference + frequentist bootstrap, used in Supp. Fig. S1 only) were inferred using RAxML-NG v1.1.0:

~~~
raxml-ng --msa <MSA> --model GTR+G4 --threads 1
~~~
- Phylogenetic trees (reference + frequentist bootstrap) were inferred using PhyML v3.3.20200621:

~~~
phyml -i <MSA> -m GTR -c 4 -a e -f e -d nt -o tlr -b 0
~~~
- Bayesian bootstrap trees were inferred using PhyML v3.3.20200621:

~~~
phyml -i <MSA> --weights=<WEIGHTS> -m GTR -c 4 -a e -f e -d nt -o tlr -b 0
~~~
- Bootstrap alignments were generated using goalign dev0537492:

~~~
goalign build seqboot -p -n 200 -i <MSA>
~~~
- Bayesian bootstrap weights were generated using goalign dev0537492:

~~~
goalign build weightboot -p -i <MSA> -n 200
~~~
- Frequentists bootstrap supports from non-collapsed bootstrap trees were computed using gotree devb324e73:

~~~
gotree compute support fbp -i  -b <BOOT> -o <OUT TREE WITH SUPPORTS>
~~~
- Bayesian and frequentist bootstrap supports from collapsed bootstrap trees were computed using gotree devb324e73:

~~~
gotree collapse length -i <BOOT> -l <0.1/length> |
gotree compute support fbp -i  -b --o <OUT TREE WITH SUPPORTS>
~~~

### Simulated datasets

- The EBOV tree from the previous dataset was used as the model tree for simulations, along with the substitution model parameters inferred by PhyML.
- Short branches (length <0.5/18,996) of this tree were replaced by random (drawn from an exponential distribution with mean 0.5/18,996) branch lengths using gotree devb324e73:

~~~
gotree brlen setrand -i <TREE> --min-len 0 --max-len 2.632133e-05 --mean 2.632133e-05
~~~
- Branches of this tree were re-scaled by different factors (X1, X4, X16, and X64) and the tree was rooted at midpoint using gotree devb324e73:

~~~
gotree brlen scale -i tmp -f <1,4,16,64> | gotree reroot midpoint
~~~
- The consensus sequence was computed as root sequence from the initial alignment using goalign devb034b00:

~~~
goalign consensus -i <MSA>
~~~
- A fraction of the sites of this consensus sequence was extracted randomly for the simulations. The number of sites depended on the scale: X1: 18996 (= EBOV MSA length); X4: 4749 = 18996/4; X16: 1187 ≈ 18996/16; X64: 296 ≈ 18996/64. We used goalign dev0537492:

~~~
goalign sample sites -l <LENGTH> --consecutive=false
~~~
- A MSA was simulated from the root-sequence and the rescaled tree using our simulator SNAG 835c479 (https://github.com/fredericlemoine/snag):

~~~
snag -model gtr -parameters 0.09084, 0.72307, 0.07212, 0.02603, 0.97950, 0.00979, 0.31900, 0.21423, 0.19823, 0.26854 -gamma -gamma- cat 4 -alpha 0.244 -intree <TREE> -root-seq <CONSENSUS>
~~~
- Reference and bootstrap trees were inferred as for the previous datasets.
- The frequentist and Bayesian supports were computed as for the previous datasets.

### SARS-CoV-2 dataset

- SARS-CoV-2 sequence identifiers were taken from Github:
  - Repository: http://github.com/evolbioinfo/phylocovid
  - File: CRAS/data/20200425/alignments/duplicates.txt
- From this file, 800 identifiers were randomly sampled:

~~~
shuf -n 800
~~~
- Sequences were extracted from GISAID archive (date: 2022_03_09) using a dedicated workflow (available on Github), and aligned using nextalign 2.7.0.
- Phylogenetically noisy positions were masked from this alignment using goalign dev0537492:

~~~
goalign mask -s 0 -l 55 -i <MSA> |
goalign mask -s 29803 -l 101 |
goalign mask --pos 186, 1058, 2093, 3036, 3129, 6989, 8021, 10322,
10740, 11073, 13407, 14785, 19683, 20147, 21136, 24033, 24377,
25562, 26143, 26460, 26680, 28076, 28825, 28853, 29699
~~~
- Reference and bootstrap trees were inferred as for the previous datasets
- The frequentist and Bayesian supports were computed as for the previous datasets.

### RVFV dataset

- RVFV sequences were retrieved from Genbank on April 17th 2024
- The sequence annotations were parsed, and sequences categorized by segment using a dedicated workflow (available on Github).
- Sequences having more than 60% gaps were removed using goalign dev0537492:

~~~
goalign clean seqs --cutoff 0.6 -i <MSA>
~~~
- For each segment, reference and bootstrap trees were inferred as for the previous datasets
- For each segment, the frequentist and Bayesian supports were computed as for the previous datasets.

### Non-virus datasets

- TreeBASE datasets were downloaded from: https://cme.h-its.org/exelixis/material/raxml_adaptive_data.tar.gz
- Alignment lengths and number of sequences were extracted using goalign dev0537492:
  - `goalign stats nseq -p -i <MSA>`
  - `goalign stats length -p -i <MSA>`
- The 300 largest DNA datasets (in number of sequences) were analyzed like the previous datasets (reference and bootstrap trees).
- The 100 largest proteic datasets (in number of sequences) were analyzed (reference and bootstrap trees) using PhyML v3.3.20200621:

~~~
phyml -i <MSA> -m LG -c 4 -d aa -a e -o tlr -b 0
~~~
- The frequentist and Bayesian supports were computed as for the previous datasets.
- From the 300 large DNA datasets, we extracted the largest MSA (in number of sites and unique sequences) with a homoplasy level <50% (sample ID 10117_0; *Psiguria* species, which belongs to Cucurbitaceae, flowering plants; see Supp. Table 1 for details, and Fig. 2 and 4 for results).

### Availability: datasets, analysis workflows and pipeline

- All our (non SARS-CoV-2) datasets and workflows are available from: https://github.com/evolbioinfo/bayesian_bootstrap_analyses/.
- SARS-CoV-2 Data Availability (796 sequences):
  - GISAID Identifier: EPI_SET_240621ps
  - doi: 10.55876/gis8.240621ps
  - All genome sequences and associated metadata in this dataset are published in GISAID’s EpiCoV database. To view the contributors of each individual sequence with details such as accession number, Virus name, Collection date, Originating Lab and Submitting Lab and the list of Authors, visit 10.55876/gis8.240621ps.
- Our pipeline BBOOT to obtain the Bayesian bootstrap supports uses PhyML. It is available at https://github.com/evolbioinfo/bayesian_bootstrap. As input, it takes a multiple sequence alignment (MSA), generates Bayesian bootstrap weights, infers reference and bootstrap trees using PhyML, and compute Bayesian supports. In addition, it produces a file containing several metrics and outputs: (1) MSA alphabet (nucleotides or proteins); (3) MSA length; (2) Minimal number of mutations in the MSA (see text); (4) Reference tree size (sum of branch lengths, in number of mutations); (5) Level of homoplasy (see text). Finally, it produces the figure showing the cumulative distribution of branch lengths (as in Fig. 2). To execute the workflow, java and singularity (or Docker) are needed to install Nextflow. Then the workflow is executed by running the following command:

~~~
nextflow run main.nf --msa <MSA>
      --results <OUT DIR: results>
      --nboot <# BOOT REPLICATES: 200>
      --collapse <COLLAPSE THRESHOLD: 0.1>
~~~

**Supplementary Table S1:**
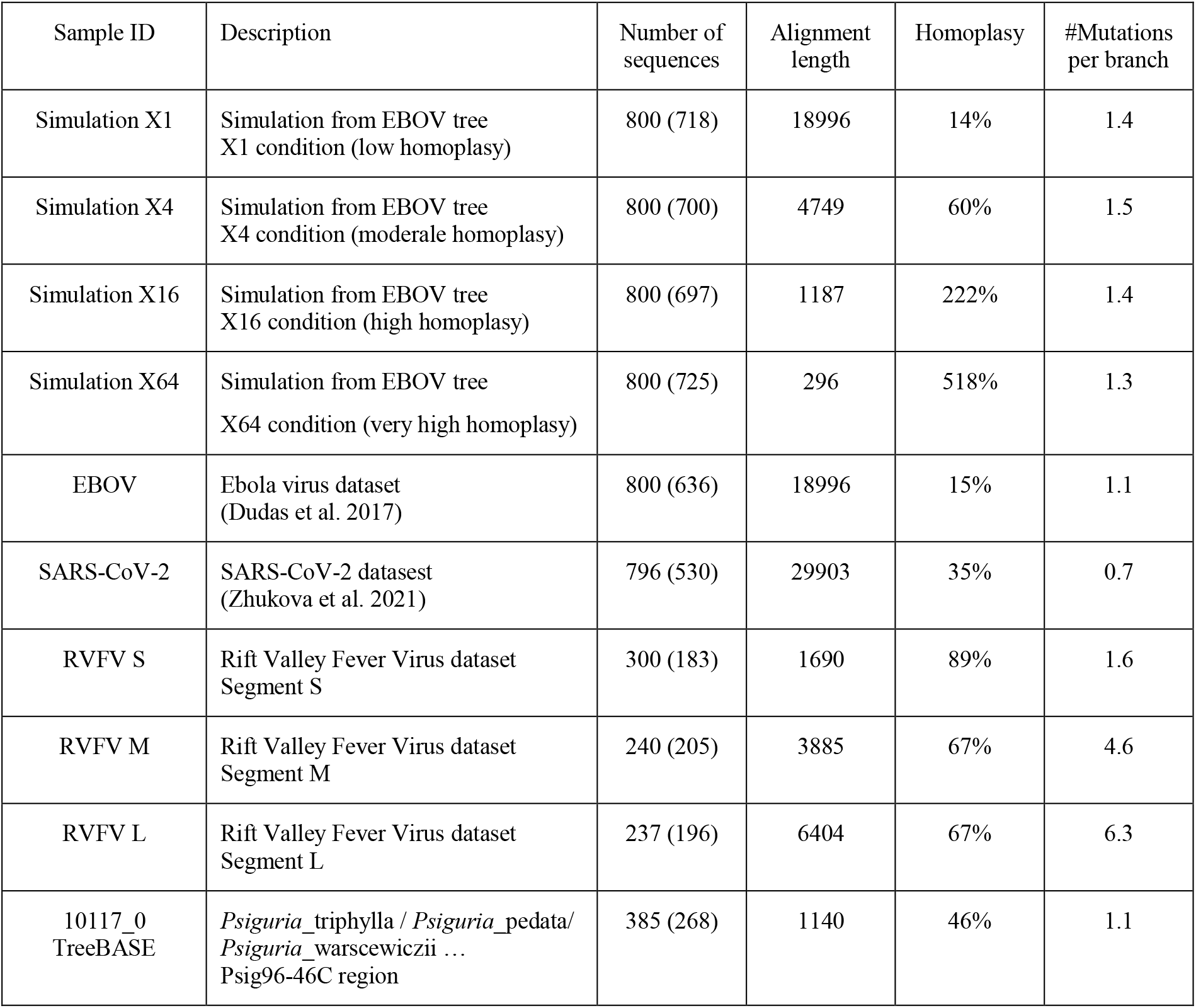
Datasets. Number of sequences: total number of sequences in the datasets, the number of unique sequences is given in brackets. Alignment length: number of sites in the MSA. Homoplasy: The level of homoplasy in the data set; assuming a perfect phylogeny, homoplasy is 0% (see text). #Mutations per branch: estimated number of mutations per branch, which is equal to the tree length (estimated by PhyML) multiplied by the MSA length and divided by the number of branches in the tree.

**Supp. Figure S1:**
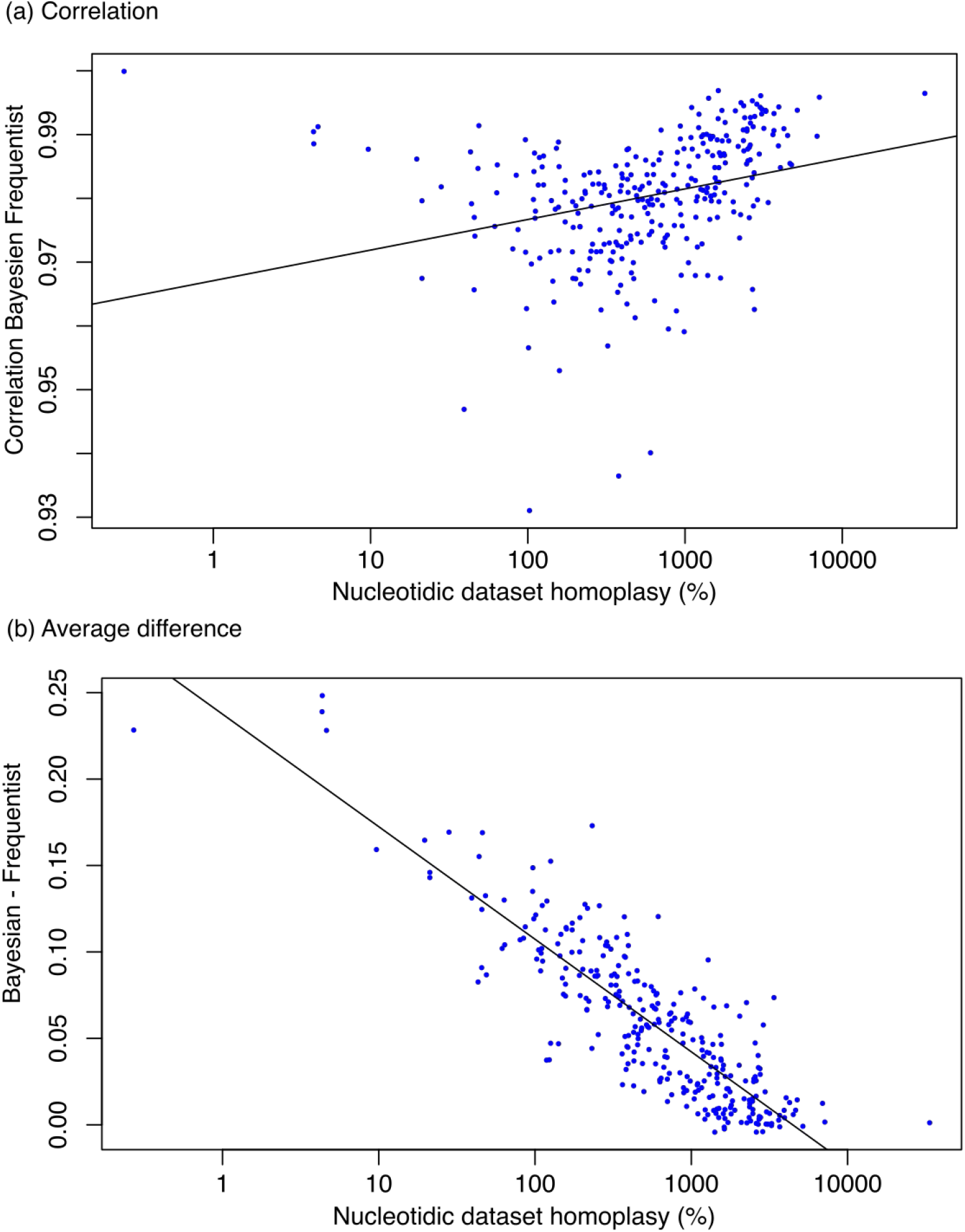
Comparison of frequentist and Bayesian supports with 300 large DNA datasets. Datasets were downloaded from: https://cme.h-its.org/exelixis/material/raxml_adaptive_data.tar.gz. The 300 largest (in number of sequences) were analyzed using the same programs and options as for the viral and non-viral low-homoplasy datasets of Supp. Table 1. **(a)** Pearson correlation between the two supports, for each of the 300 datasets. **(b)** Average difference between the Bayesian and CollapseFrequentist supports, for each of the 300 datasets (excluding branches with near-zero length, for which both supports are close to zero). The average homoplasy level is 1,210%, but 17 datasets have less than 50% homoplasy; the plant dataset of Figures 2 and 4 is the largest of these 17, with homoplasy equal to 46%. **(a)** The Pearson correlation between the two supports is always high (average: 98%) and increases with the level of homoplasy, as expected, since both supports become almost equal with high homoplasy. **(b)** We observe that the average difference between the supports is quite high at low homoplasy levels (up to 25%) and then decreases to be close to zero, as expected.

**Supp. Figure S2:**
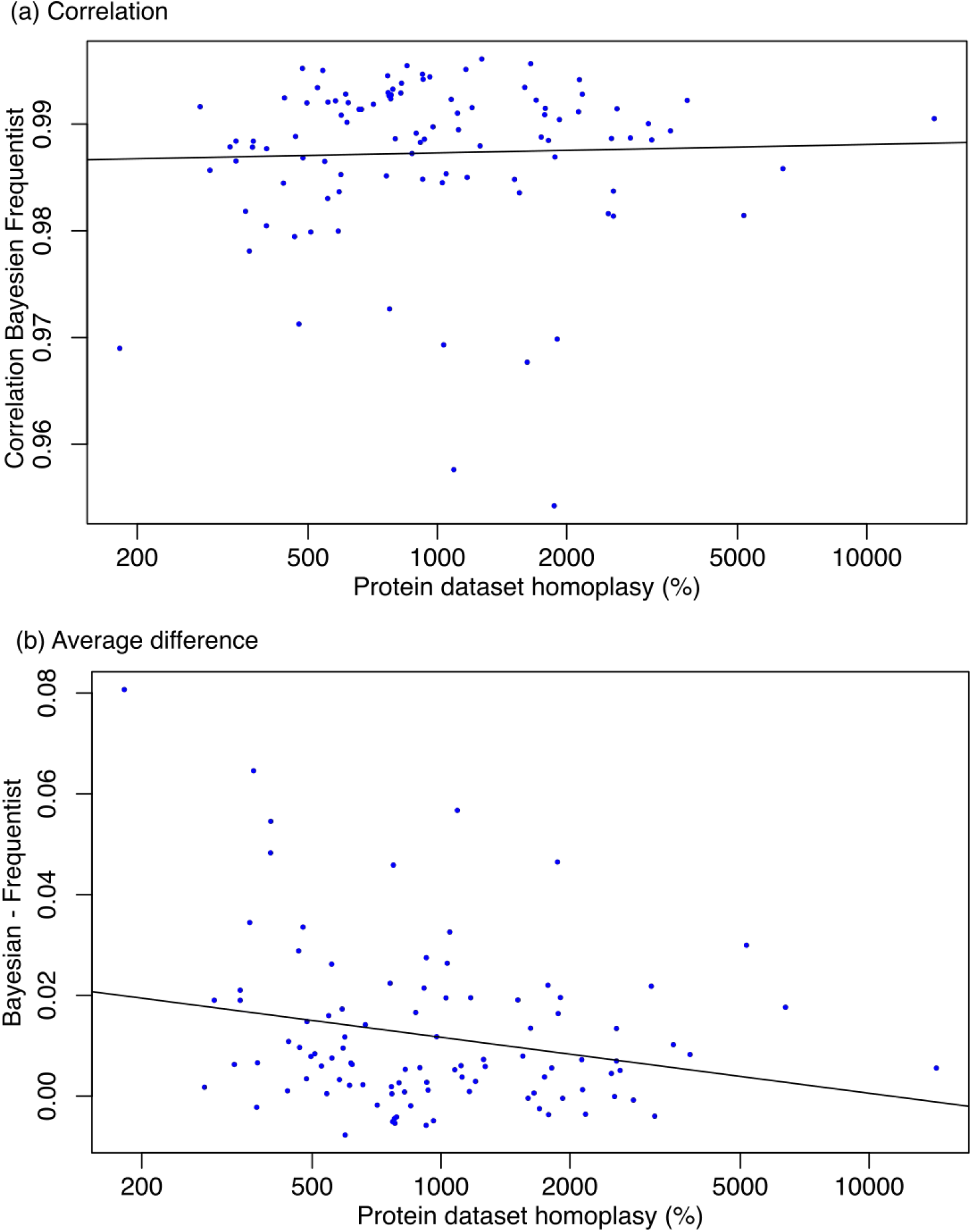
Comparison of frequentist and Bayesian supports with 100 large protein datasets. Datasets were downloaded from: https://cme.h-its.org/exelixis/material/raxml_adaptive_data.tar.gz. The 100 largest (in number of sequences) were analyzed using the same programs and options as for the viral and non-viral low-homoplasy datasets of Supp. Table 1, but using substitution model LG+G4. **(a)** Pearson correlation between the two supports, for each of the 100 datasets. **(b)** Average difference between the Bayesian and FrequentistCcollapse supports, for each of the 100 datasets (excluding branches with near-zero length, for which both supports are close to zero). The average homoplasy level is 1,351%, with minimum homoplasy equal to 182%, i.e. much higher than with DNA datasets, probably because protein datasets in TreeBASE were generally used to elucidate deep branchings. As with DNA, we observe that the gap between the two supports narrows with high homoplasy, and that the correlation between the two supports is strong for all datasets (>95%, 99% on average).

